# Multilevel Twin Models: Geographical Region as a Third Level Variable

**DOI:** 10.1101/2020.11.11.377820

**Authors:** Zenab Tamimy, Sofieke T. Kevenaar, Jouke Jan Hottenga, Michael D. Hunter, Eveline L. de Zeeuw, Michael C. Neale, Catharina E.M. van Beijsterveldt, Conor V. Dolan, Elsje van Bergen, Dorret I. Boomsma

## Abstract

The classical twin model can be reparametrized as an equivalent multilevel model. The multilevel parameterization has underexplored advantages, such as the possibility to include higher-level clustering variables in which lower levels are nested. When this higher-level clustering is not modeled, its variance is captured by the common environmental variance component. In this paper we illustrate the application of a 3-level multilevel model to twin data by analyzing the regional clustering of 7-year-old children’s height in the Netherlands. Our findings show that 1.8%, of the phenotypic variance in children’s height is attributable to regional clustering, which is 7% of the variance explained by between-family or common environmental components. Since regional clustering may represent ancestry, we also investigate the effect of region after correcting for genetic principal components, in a subsample of participants with genome-wide SNP data. After correction, region did no longer explain variation in height. Our results suggest that the phenotypic variance explained by region actually represent ancestry effects on height.

## Introduction

The classical twin model (CTM) is often approached from a structural equation modeling (SEM) framework (Bentler and Stein, 1992; Boomsma and Molenaar, 1986; Heath et al., 1989; Neale & Cardon, 1992; Rijsdijk and Sham, 2002). In this framework, it is a one-level model with family as level one sampling unit. The analysis of twin data can, however, also be approached from a multilevel model (MLM) perspective. MLMs were developed specifically for the analysis of clustered data (Goldstein, 2011; Laird and Ware, 1982; Longford, 1993; Paterson and Goldstein, 1991). Classical examples are children (level 1 units), who are clustered in classes (level 2) within schools (level 3; Sellström and Bremberg, 2006). Other examples are fMRI measures (level 1) that are clustered in individuals (level 2), who are clustered in scanner type (level 3; Chen et al., 2012), or biomarker data (level 1) that are clustered in measurement batches (level 2; Scharpf et al., 2011). The classical twin design is based on data that also have natural clustering, namely, twins are clustered within pairs. For this reason, the CTM can be accommodated in the MLM framework (Guo and Wang, 2002; McArdle and Prescott, 2005; Rabe-Hesketh et al., 2008; Van den Oord, 2001;). Hunter (this issue) provides a detailed account of the CTM in the MLM framework with example code and several extensions. While the MLM specification of the CTM is equivalent to the SEM approach, it also has some interesting, yet underexplored, advantages. In this paper we aim to elaborate on these advantages, and to provide an empirical illustration of a multilevel twin model, where we study the clustering of children’s height in geographical regions in the Netherlands, and consider the role therein of genetic ancestry.

In the SEM approach to the CTM, the covariance structure of twin-pairs is modelled to decompose phenotypic variance into multiple components that represent genetic and non-genetic influences. Given the biometrical underpinning of the twin model (Eaves et al. 1978; Falconer and MacKay, 1996; Fisher, 1918), the phenotypic variance can be decomposed into additive genetic variance (A), non-additive or dominance genetic variance (D) or common environmental variance (C), and unique environmental (E) variance components. Variance decomposition is based on the premise that monozygotic (MZ) twins share 100% of their DNA and dizygotic (DZ) twins share on average 50% of their segregating genes. Hence, additive and non-additive genetic variance is fully shared by MZ twins, whereas additive and non-additive variance components are shared for 50 and 25% by DZ twins. In the CTM, all influences that are not captured by segregating autosomal genetic variants are labeled as “environment”. These influences can be distinguished into common environment (shared by twins from the same family) and unique environment (creating variation among members from the same family) and are also referred to as between and within family influences. The full ACDE model is not identified when analyzing one phenotype per twin, and therefore only three of the four components can be simultaneously estimated. In this SEM approach to modeling twin data, the variance decomposition is based on the bivariate data observed in twin pairs (i.e., one phenotype for twin 1, and one for twin 2, which are both level 1 units).

In the MLM framework the phenotypic variance can be decomposed into a within-pair (level 1) and a between-pair (or family; level 2) components. This requires reparameterization of the model into level 1 and level 2 variance components. Because the E component captures variance that is not shared by twins, this component is an individual level 1 variance component. The C component is by definition shared by twins, regardless of zygosity, and is a family level 2 variance component. The A component, however, is more complicated, as it is a level 2 component in MZ twin pairs, but both a level 1 and a level 2 component in DZ twin pairs. To account for this, the A-component is divided into two orthogonal components, unique additive (A_U_) and common additive (A_C_). Here, A_U_ is a first-level component representing the A variance at the individual level (within pairs or within families), while A_C_ is a second-level component (between pairs or between families), representing the A variance at the twin-pair level. These definitions are consistent with the classical notations in which A_C_ refers to within family genetic variance known as A_1_ (Boomsma and Molenaar, 1986; Martin and Eaves 1977), or the average breeding value variance (Barton et al., 2017), while A_U_ refers to the between family genetic variance known as A_2_ (Boomsma and Molenaar, 1986; Martin & Eaves 1977), or the segregating genetic variance (Barton et al., 2017). In MZs, the A_U_ variance component is 0, since all the variance explained by A is shared by both twins from a pair. For DZ twins, the variance of both A_C_ and A_U_ are constrained to equal 0.5, since on average 50% of the A variance is shared by the individuals and 50% of the A variance is unique for the individual.

An important, but yet underexplored, advantage of the MLM approach, is the possibility to include higher-level variables in which lower-levels are nested. By including these higher-level variables, we can identify variance components which are attributable to higher-level clustering. Such clusters may be a consequence of data acquisition or design, e.g., clustering of biomarker data that are measured in batches, or clustering of brain imaging data by fMRI scanner type. They may also occur naturally, for example, families in regions, neighborhoods or schools. If the higher-level variable is not included in the variance decomposition models, the variance that it explains will be captured as part of the C-component, since both twins, regardless of zygosity, share the higher-level variable (i.e., the twin pair is nested in the higher-level variable).

Within the SEM framework, higher-level variables can be included in the model as a fixed effect on the individual level (i.e., covariate) by means of (linear) regression. For covariates (a.k.a. factors in the ANOVA sense) that do not have a linear relation with the outcome variables, this approach requires the variable to be dummy coded, which may be impractical, for example when the number of assays for a biomarker or the number of schools that twins are enrolled in, is large. In the MLM framework, however, the higher-level variable is treated as a random rather than a fixed effect, and this reduces the number of parameters to one single variance component, albeit subject to assumptions That is, given a factor with L categories, the fixed effects approach requires L-1 parameters, while the random effects approach requires one parameter (a variance component). In addition, the MLM approach is more suitable to deal with unequal group sizes (Gelman, 2005). Finally, an MLM approach allows us to evaluate the contribution of the higher-level component to the C-component, as estimated in the standard twin model. This can be achieved by comparing the C-component estimate of the two-level model (i.e., the standard twin mode) to the estimate of the three-level model.

In this paper, we illustrate the use of multilevel twin models by investigating the regional clustering of children’s height with twin data from the Netherlands Twin Register (Boomsma et al., 1992; Ligthart et al., 2019). Height serves as an indicator of the general development of a country, and is known to decrease in times of scarcity and increase in times of prosperity (Baten & Blum, 2014; Baten & Komlos, 1998). Also, children’s height is an indicator of overall development, where height is associated with cognitive development and school achievement (Karp et al., 1992; Spears, 2012). In 7-year-old children, resemblance between family members for height is explained by additive genetic (approximately 60%) and common environmental (approximately 20%) factors (Jelenkovic et al., 2016; Silventoinen et al., 2004; Silventoinen et al., 2007).

In the Netherlands there are well-established associations between height and geographical region (Abdellaoui et al. 2013), which makes this a clustering variable of interest. Geographical region will represent genetic and environmental differences between inhabitants. Location is associated with genetic differences (e.g. Colodro-Conde et al. 2018) and differences in social and cultural traditions, diet, socio-economic status, and living circumstances (e.g., rural vs urban). By analyzing height and geographical region data in a three-level MLM, we can uncover whether variation in children’s height is associated with geographic region, and identify the proportion of the common environmental or between-family variance that can be explained by these regional effects.

In a subsample of 7-year-old participants, we next investigate the extent to which regional clustering may be due to genetic ancestry by including the first three genetic principal components (PCs; Hotelling, 1933). The genetic PCs are obtained through principal component analysis of the covariance matrix of the genotype (Single Nucleotide Polymorphism (SNP) data (Reich et al., 2008). In the Netherlands, the first genetic PC is associated with a north-south height gradient (Abdellaoui et al., 2013; Boomsma et al., 2014). This gradient represents social, geographical and historical divisions between the north and the south. Southern regions were conquered by the Roman empire, adopted Catholicism, and were geographically separated from the northern regions by the waterline of the five large rivers in the Netherlands (Schalekamp, 2009). The second PC is associated with the east-west division of the Netherlands and reflects differences between rural and urban environments, since the east of the Netherlands is characterized by less populous and rural areas, while the west includes the Randstad, which is the largest urban metropole of the country. The third PC is associated with the more central regions of the country (Abdelloui et al. 2013). By adding the PCs to our models, we assessed the role of genetic ancestry of individuals between regions.

The outline of this paper is as follows. We first considered regional clustering of children’s height in a large data set of MZ and DZ twins (N = 7436). Next, we considered the model within a subgroup of children who were genotyped on genome-wide SNP arrays (N = 1375). Subsequently, we determined whether the region effects represent genetic ancestry. To this end, we analyze the relationship between the three PCs and height in 7-year-old children, and include the genetic PCs that show an association as an individual level (level 1) covariate in the model.

## Methods

### Participants and procedure

The data were obtained from the Netherlands Twin Register (NTR), which has collected data on multiple-births and their family members since 1987 (Ligthart et al., 2019). In the longitudinal NTR surveys of phenotypes in children, parents were asked to complete questionnaires on their children’s health, growth and behavior with intervals of approximately two years.

For the present study, we included data on 6- and 7-year-old twin children (range 6 years and 0 months – 7 years and 11 months). The analyzed sample included 7,346 twin children (50.3% girls), in 3724 families. The twins were 7.4 (SD 0.3) years old on average, when their mothers reported their height. Of these children, 1,375 (18.7% of total) were genotyped. Genotyping largely took place independent of phenotype criteria. The 1,375 genotyped individuals were from 714 families, 52.4% of this subsample were girls and the average age was 7.4 (SD 0.3). In this study, we included participants with both height and postal code information at age 7. Mothers were asked to report the four digits of the current postal code in the questionnaire at age 7 since 2002. This study includes data collected between 2002 and 2015, with children from birth cohorts 1995 through 2007. Postal code data were missing for approximately 1% of the surveys that were sent out after 2002, this may reflect participants whose parents had for example moved abroad. Approximately 20% of the parents did not report their children’s height at age 7. A flowchart outlining the sample size after every step of exclusion is displayed in Figure 1. Only twins were included in the initial sample; singletons or triplets were excluded. Children with severe handicaps were excluded, as were multiple twin pairs per family, twins born before 34 weeks of gestation, and twins outside the 6-8 age range. Zygosity was determined by DNA polymorphisms or by a parent-reported zygosity questionnaire on twin similarity. The zygosity determination by questionnaire has an accuracy of over 95% (Ligthart et al., 2019). Table I displays the descriptive statistics of the phenotypic data by zygosity for the total and for the genotyped sample.

**Table I.**
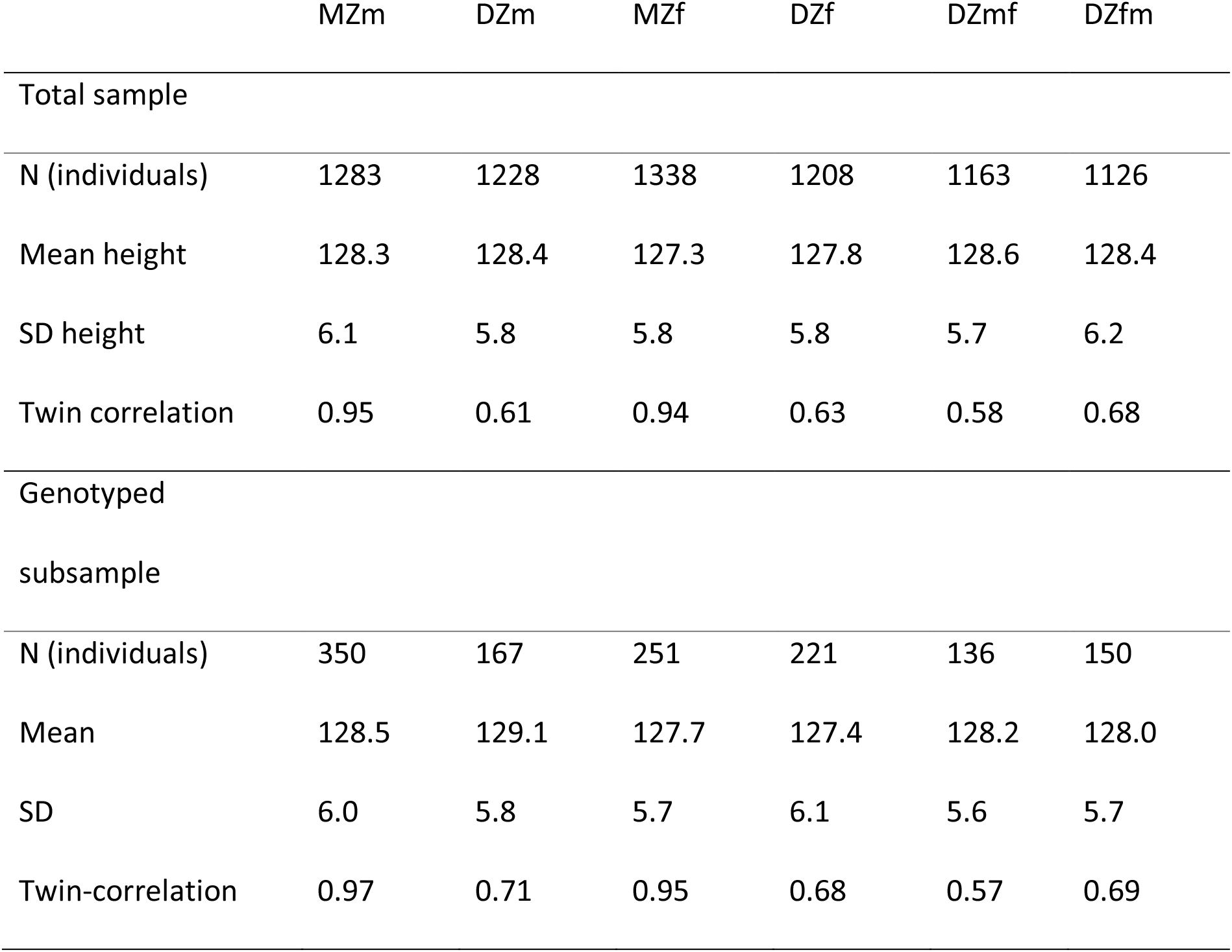
The number of twins, the mean, standard deviation and the twin correlation per zygosity group for the total sample and the genotyped subsample.

**Figure 1.**
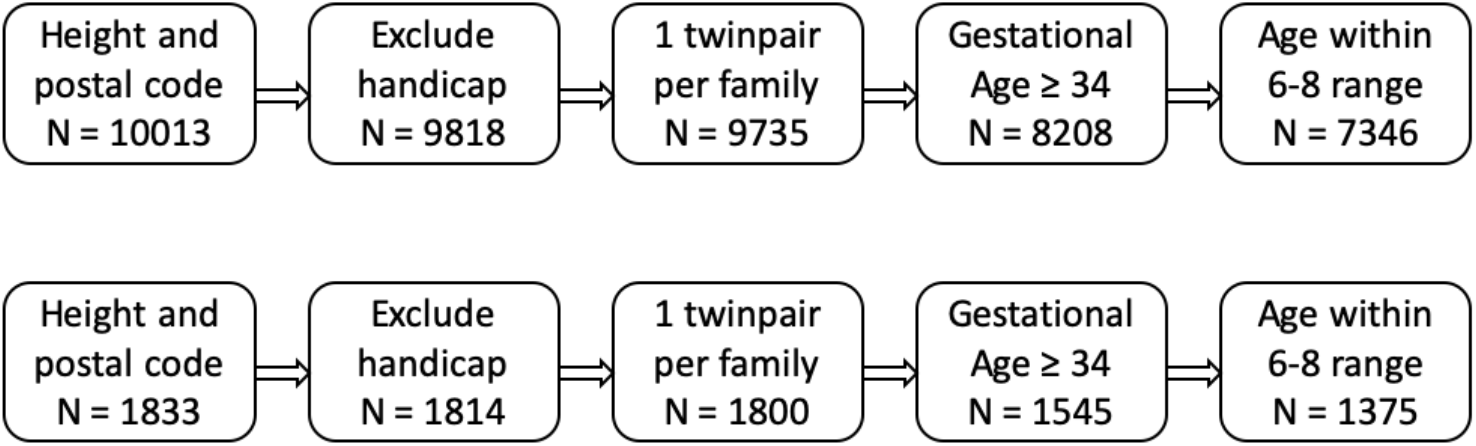
Flowchart containing sample size for the total sample (upper row) and the genotyped subsample (lower row) after every step of exclusion.

### Measures

#### Height

Height in centimeters was reported by mothers. In the questionnaire, mothers reported the height of the child as well as the date of the measurement. Maternal report of height correlates .96 with height measured in the laboratory (Estourgie-van Burk et al., 2006). Mothers reported the age of their children at the moment of completing the survey and the date of the height measurement. For 5% of the children, the date at the time of height measurement was not available. Therefore, in this 5%, we took the age at the time of questionnaire completion. The correlation between age at questionnaire completion and age at height measurement is 0.95, and the mean difference in age is 0.01 years.

#### Region

At the time of reporting height, parents also reported the four digits of the postal code of their current address. In the Netherlands, postal codes map to geographical locations. The postal code consists of four digits and two letters, where the first two digits map to region and the second two digits and letters map to city, neighborhood within the city, and street. In our analyses, region is specified by the first two digits of the postal code, resulting in 90 regions which are displayed in Figure 2. They cover on average 462 km^2^ and have a mean population of around 192,000 (total for the country is 41,543 km^2^, including ~19% water). Most regions encompass several municipalities. In the total sample, the number of children per postal code unit ranged from 10 to 194 (M= 81.6, SD = 38.4). In the genotyped sample, the number of children per postal code unit ranged from 1 to 43 (M= 15.6, SD = 8.6).

**Figure 2.**
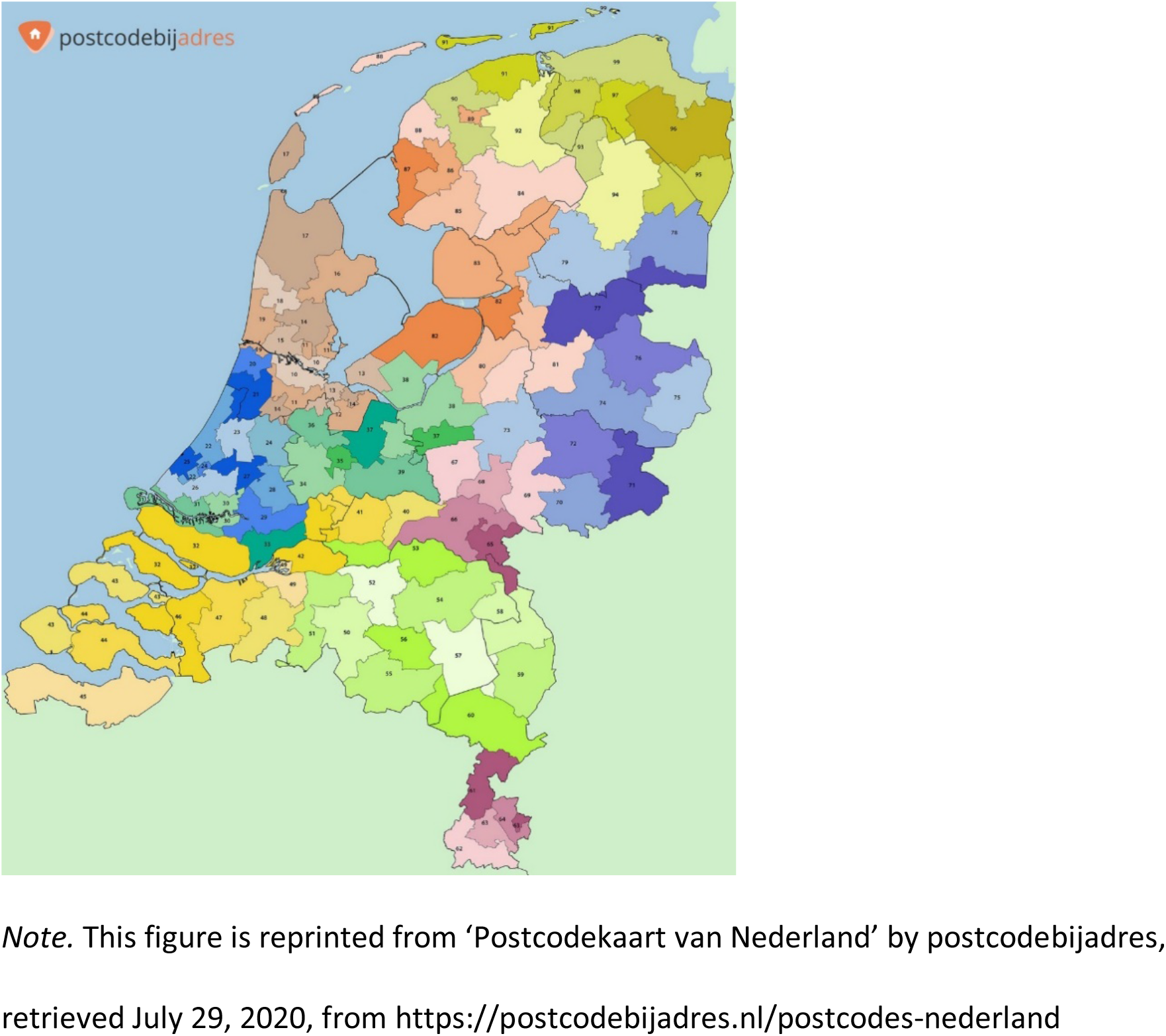
Map of the 90 regions in the Netherlands based on first two digits of the postal code. *Note.* This figure is reprinted from ‘Postcodekaart van Nederland’ by postcodebijadres, retrieved July 29, 2020, from https://postcodebijadres.nl/postcodes-nederland

#### Principal components

Genotype data in 1375 individuals were collected by the following genotype platforms: Affymetrix 6, Axiom and Perlegen, Illumina 1M, 660 and GSA-NTR. The SNP data obtained on the 6 platforms were pruned in Plink to be independent, with additional filters to ensure Minor Allele Frequency (MAF) > 0.01, Hardy-Weinberg Equilibrium (HWE) p > 0.0001 and call rate over 95%. Subsequently, long range Linkage Disequilibrium (LD) regions were excluded as described in Abdellaoui et al. (2013), because elevated levels of LD result in overrepresentation of these loci in the PCs, disguising genome-wide patterns that reflect ancestry. For each platform, the NTR data were merged with the data of the individuals from the 1000 Genomes reference panel for the same SNPs, and Principal Components were calculated using SMARTPCA (Prince et al., 2006), where the 1000 genomes populations were projected onto the NTR participants (Abdellaoui et al., 2013). Population outliers were identified using pairwise PC plots and excluded, rendering the final clustering homogeneous. The NTR platform genotype data of this cluster were aligned to the GoNL reference panel V4, merged into a single dataset, and then imputed using MachAdmix. From the imputed data, SNPs were selected that satisfied R^2^ ≥ 0.90, and that were genotyped on at least one platform. These SNPs were subsequently filtered on MAF < 0.025, HWE p < 0.0001, call rate ≥ 98%, and the absence of Mendelian errors. Again, the long-range LD regions were removed from these SNP data. With this selection of SNPs, 20 new PCs were calculated with SMARTPCA (Prince et al., 2006), to indicate the residual Dutch genetic stratification.

### Models

#### The Classical Twin Model

In the classical twin model, the phenotypic variance can be decomposed into three components: Additive genetic (A), Common environmental (C) and unique Environment (E) component, which includes measurement error. As in most earlier publications, we will not consider genetic dominance variance for height (but see Joshi et al., 2015). The variance component model, with subscript *i* for individuals, is:

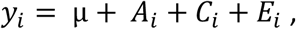

where μ is the intercept (phenotype mean), and A_i_ ~ N(0, σ^2^_A_) is the additive genetic deviation, *C_i_* ~ N(0, σ^2^_C_) is the common environmental deviation, and *E_i_* ~ N(0, σ^2^_E_) is the unique environmental deviation.

Assuming A, C, and E are mutually independent, we have the following decomposition of phenotypic variance:

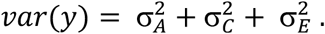

Alternatively, the variance component model can be written as a path model in which A, C and E are standardized to have unit variance (see Figure 3):

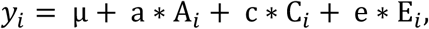

**Figure 3.**
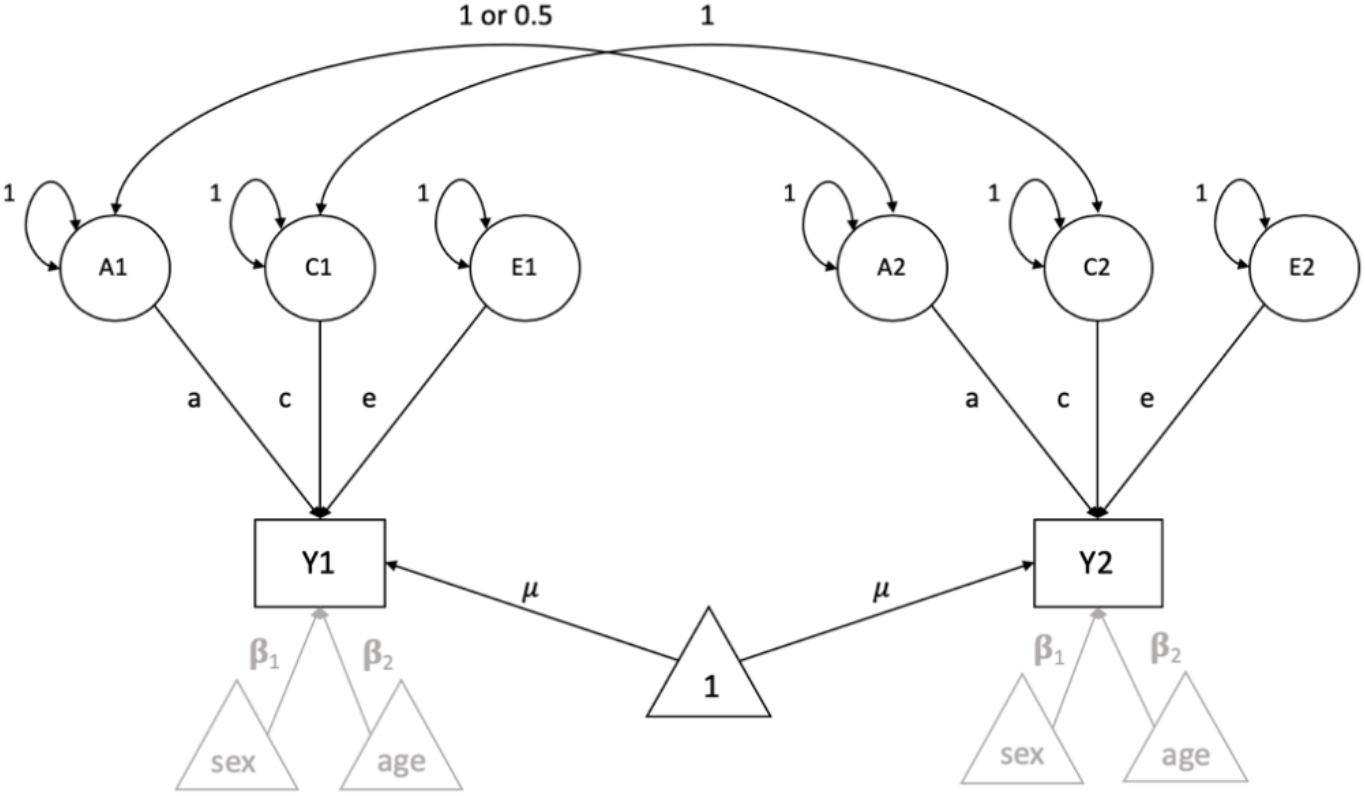
The Classical Twin Model including three latent factors per person, representing Additive genetic, Common and unique Environmental influences. Two additional covariates, age and sex, are presented in a schematic way in grey.

Here, the variance decomposition is:

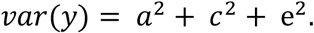

In terms of the path coefficient model, the covariance between the twins equals σ_*mz*_ = *a*^2^ + *c*^2^, in MZ twins, and 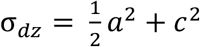 in DZ twins.

#### Multilevel Twin Model

When specifying a CTM as an MLM, the variance components of the CTM are parametrized as within- and between family components. The additive genetic variance is separated into two parts: a part that is shared by the members of a twin pair on the second level, A_C_, and a part that is unique to each individual on the first level, A_U1_ and A_U2_). The path coefficients associated with the A_C_ and A_U_ are equal. The variance of the common genetic factor (r) and the unique genetic factor (1-r) depend on the zygosity of the twin pair: for MZ r = 1.00, while for DZ r = 0.50. The common environmental factor, representing between family influences, is a level two component. Unique environmental factors E represent within family, level one, influences. The means (intercepts) μ are specified on the first level and are assumed to be equal for first- and second-born twins and zygosity. The ACE model in multilevel parametrization is illustrated in Figure 4. Here, we included age at the individual level, because it represents the age at reported height measure and could differ between twins.

**Figure 4.**
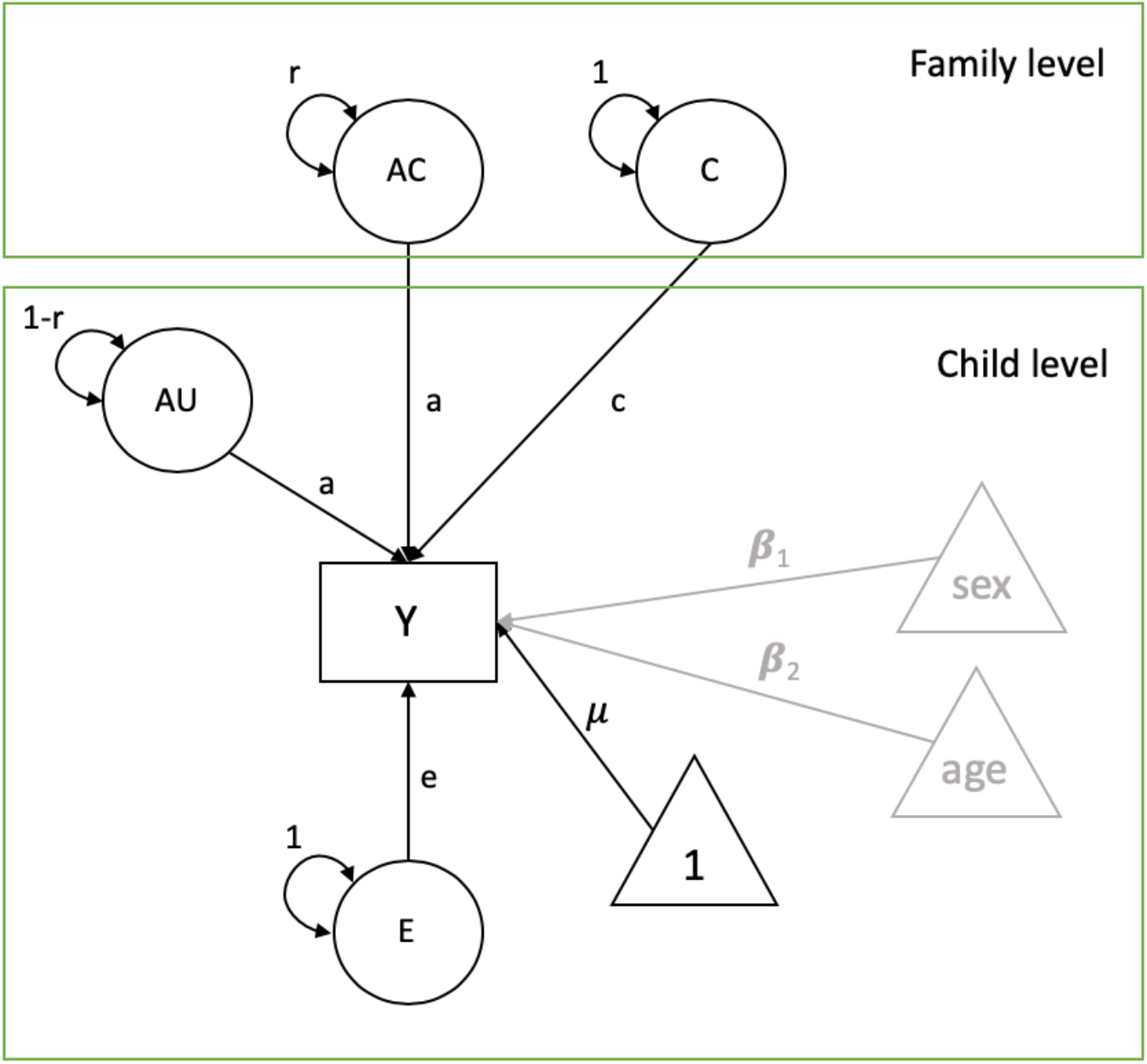
Multilevel parametrization of the ACE model, where Y represents the phenotype, latent variables A_U_ represent the unique additive genetic influences and E unique environment. μ is the intercept, sex and age are covariates, presented in a schematic way. On the family level, C is common environment and A_C_ common genetic influences. The path coefficients, a, c, e, β_1_ and β_2_ represent regression coefficients. The r parameter represents variance (1 for MZ twins and 0.5 for DZ twins).

#### Multilevel Twin Model with third level clustering variable and individual level covariates

Other clustering variables can be added to this model, as displayed in Figure 5. A higher-order clustering variable can be added to the third level of this model in two steps. On the third level, the higher-order clustering variable is added with a variance of 1 and a path loading of 1 to a latent variable on the second level, which has a variance of 0 and a freely estimated path loading from Region (reg) to the observed phenotype. The same 3-level model which also includes PC1 as a fixed covariate is displayed in Figure 6.

**Figure 5.**
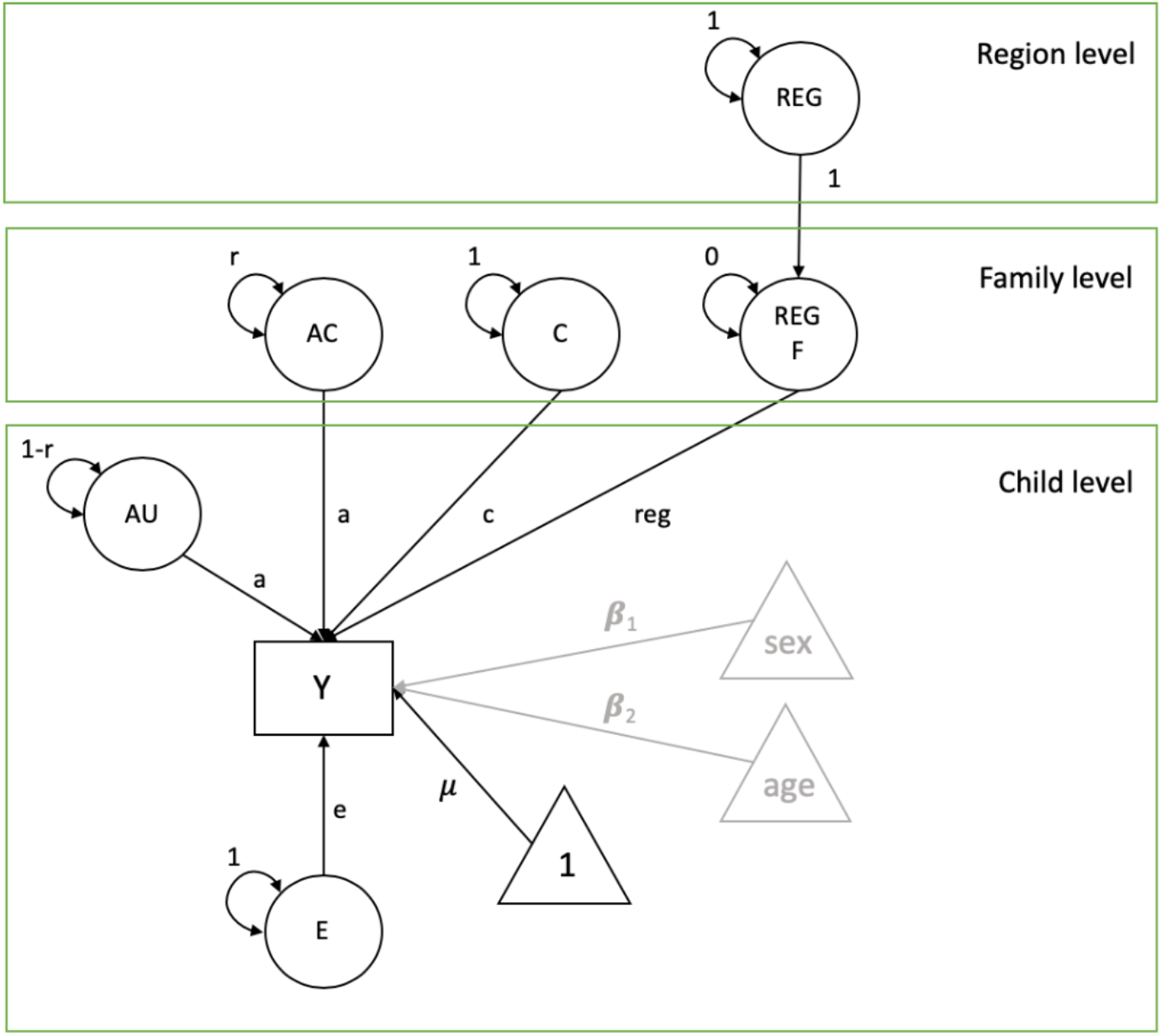
Multilevel parametrization of the ACE model with Region as a third level, which loads on the region variable REG F on the family level, on which the observed variable Y is regressed with its coefficient estimating the effect of region.

**Figure 6.**
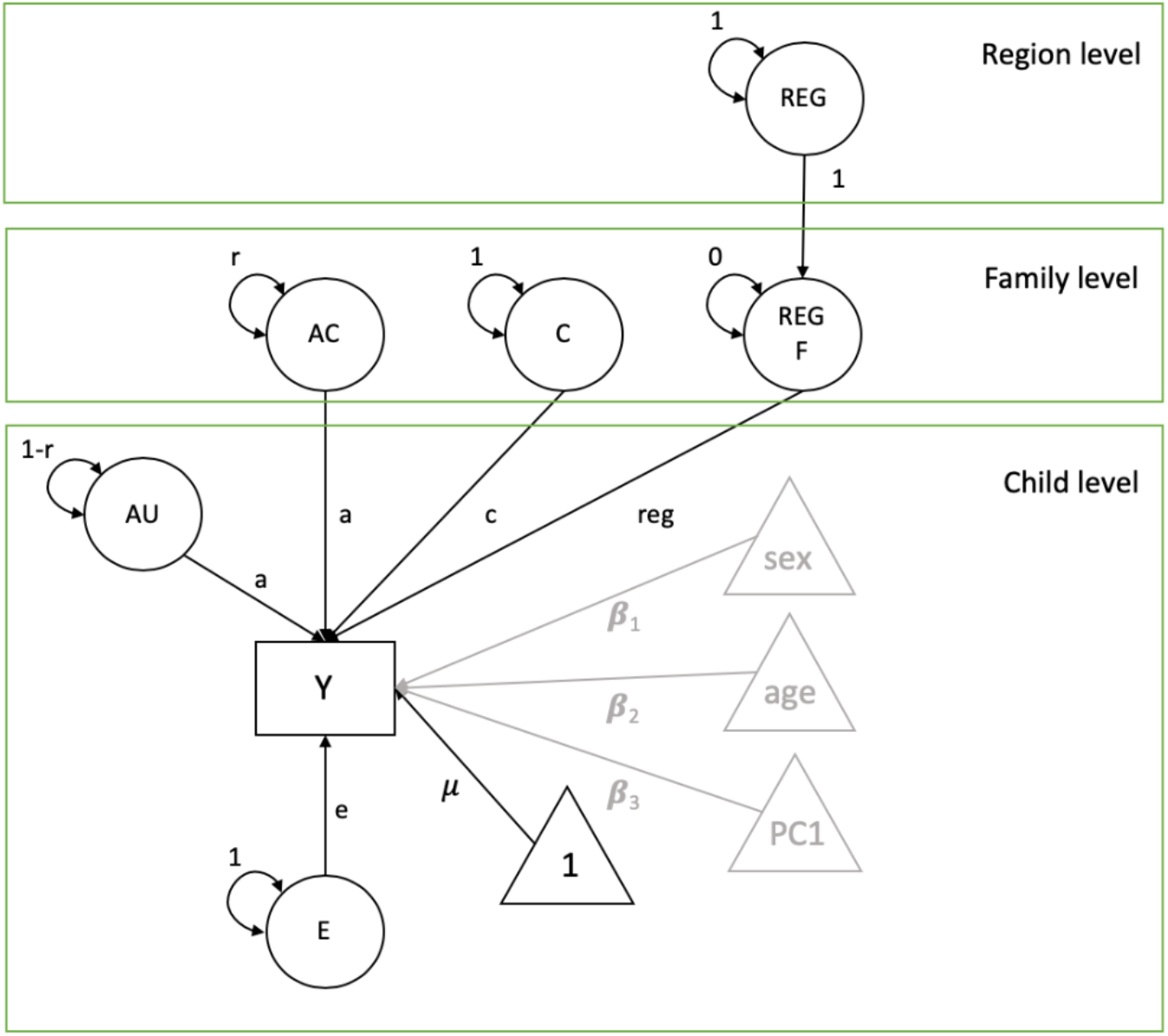
Multilevel parametrization of the ACE model with Region as a third level, and including PC1 as a covariate (we test PC1, PC2, and PC3, but depict only PC1 to avoid clutter, and because our final model included only PC1).

#### Analyses

All analyses were performed in R (R Core Team, 2020) with the package OpenMx (Boker et al., 2011; Neale at al., 2016; Pritkin et al., 2017). Age at measurement was converted to z-scores and scores on PCs were multiplied by 1000 to facilitate optimization. First, in the full sample, a variance decomposition of the variance in height was obtained in the regular CTM. We included the z-scores of age at measurement and sex as covariates. Then, we repeated the analysis in the multilevel model to illustrate the equivalence of the two approaches. Following this, we added region as a third level in the multilevel parametrization. We repeated these steps in the genotyped group to investigate the representativeness of this subsample. Finally, in the genotyped subset, we added the PC scores as individual level covariates in the 3-level model.

We tested the contribution of region to the variance of height by comparing the difference of fit in the 3-level model and the 2-level model without region with a log-likelihood ratio test. Under certain regularity conditions (Steiger et al., 1985), the difference in fit between these models is distributed as chi-squared with one degree of freedom.

## Results

When we plotted the average height data by region, a north-south trend was distinguished, with the children in the northern regions somewhat taller than those in the southern part of the Netherlands (of the 12 provinces in the Netherlands, the northern province Drenthe had the highest mean height (M = 129.40) and the southern province Noord-Brabant had the lowest mean height (M = 127.01)). Figure 7 displays the mean height of 7-year-olds per region. In the genotyped group, height correlated with PC1 (i.e., the PC showing a North-South gradient) (r = 0.16), but not with other PCs (r = 0.01 for PC2, r = −0.01 for PC3). Therefore, we incorporated PC1 into subsequent analyses and omitted PC2 and PC3.

**Figure 7.**
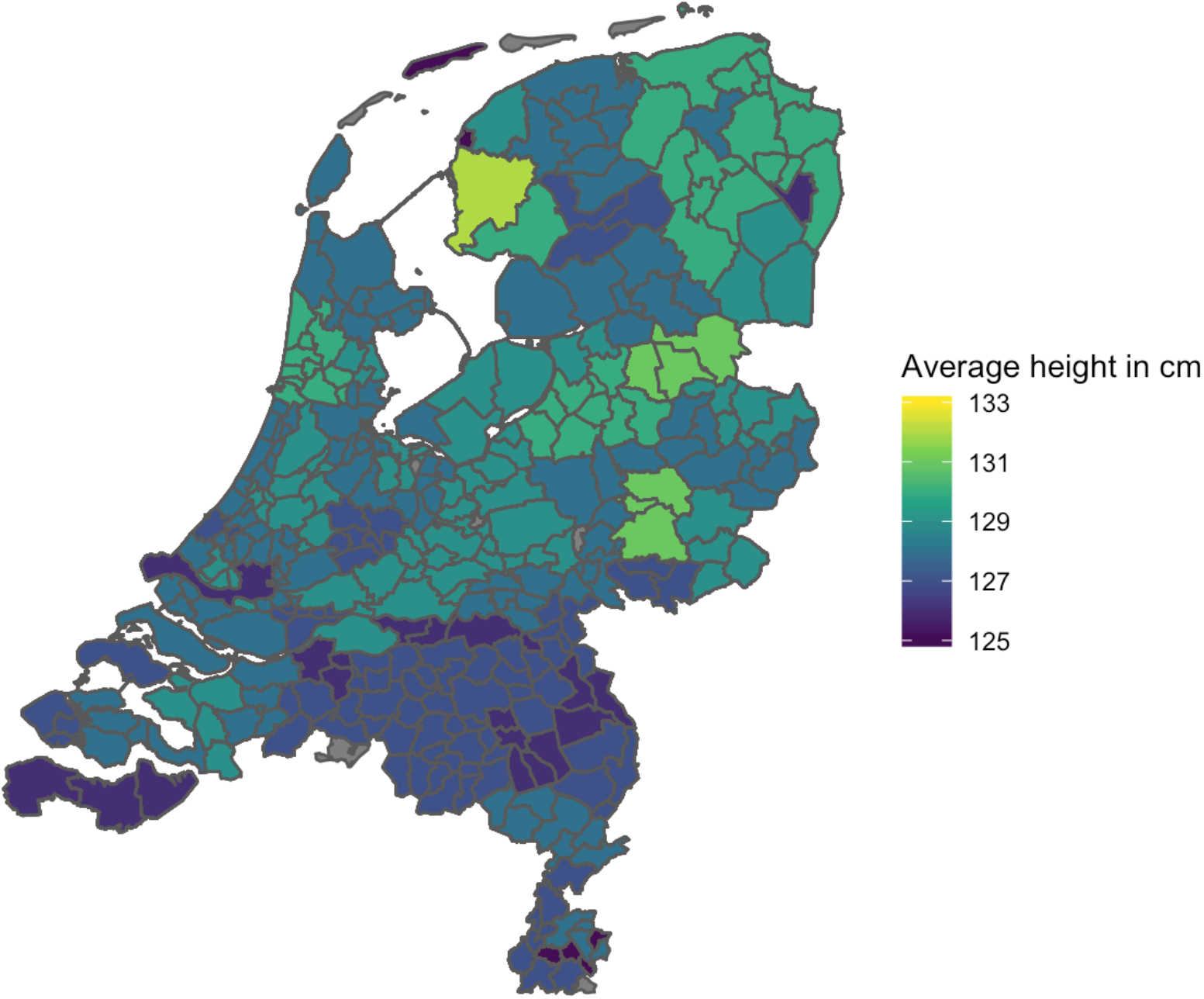
Mean height of 7-year old children in centimeters by region in the Netherlands.

The 2-level model fitted significantly worse than the 3-level model with region as level three clustering variable (Δ-2LL = 22.93, Δdf = 1, p <.001). So, region in the Netherlands associates with a statistically significant proportion (1.8%) of the variance in height in 7-year-olds. Table II displays the parameter estimates and the standardized variance components of the models. Comparing the parameter estimates of the models shows that the variance attributable to region in the 3-level model was captured by the C-component in the 2-level model.

**Table II.** Results of CTM and 2-level and 3-level MLM analyses for the full sample: path coefficient estimates with standard errors (SE) and standardized variance components of the 2-level and the 3-level models (with age and sex as covariates). N = 7346 twins in 3724 families.

|  | parameter | CTM | 2-level model | 3-level model |
| --- | --- | --- | --- | --- |
| Intercept | Intercept (SE) | 128.4 (0.11) | 128.4 (0.11) | 128.5 (0.14) |
| Covariates | $\beta_{\text{sex}}$ (SE) | -0.62 (0.13) | -0.63 (0.12) | -0.62 (0.12) |
| | $\beta_{\text{age}}$ (SE) | 1.40 (0.09) | 1.42 (0.08) | 1.42 (0.08) |
| a, c, e, region path loadings | a (SE) | 4.74 (0.10) | 4.70 (0.08) | 4.70 (0.08) |
|  | c (SE) | **2.87 (0.18)** | **2.90 (0.16)** | **2.80 (0.16)** |
|  | e (SE) | 1.39 (0.03) | 1.39 (0.03) | 1.39 (0.03)) |
|  | region (SE) |  |  | 92.33 (2.13) |
| Total variance |  | 32.70 | 32.46 | 32.47 |
| $(a^2 + c^2 + e^2 (+ \text{region}^2))$ | | | | |
| A (standardized) |  | 0.688 | 0.681 | 0.681 |
| C (standardized) |  | **0.252** | **0.259** | **0.241** |
| E (standardized) |  | 0.059 | 0.060 | 0.060 |
| REGION (standardized) |  |  |  | 0.018 |

### Results analyses genotyped sample

In the genotyped group, region explained 1.6% of the variance, which almost equals the percentage reported above. The likelihood ratio test of this component was not significant: Δ-2LL = 0.85, Δdf = 1, p = .36. However, we ascribed this to a lack of power given the appreciably smaller sample size (in terms of individuals, N = 7,346, vs. N = 1,375). The parameter estimates and standardized variance components are displayed in Table III.

**Table III.** Results of CTM and 2-level and 3-level MLM analyses in the genotyped sample (N = 1375 twins in 714 families. Path coefficient estimates with standard errors (SE) and standardized variance components of the 2- and 3-level model (with age and sex as covariates).

|  | parameter | 2-level model | 3-level model |
| --- | --- | --- | --- |
| Intercept | Intercept (SE) | 128.4 (0.26) | 128.5 (0.27) |
| Covariates | $\beta_{\text{sex}}$ (SE) | -0.62 (0.31) | -0.63 (0.31) |
| | $\beta_{\text{age}}$ (SE) | 1.16 (0.18) | 1.15 (0.18) |
| a, c, e, region path loadings | a (SE) | 4.53 (0.20) | 4.53 (0.20) |
|  | c (SE) | **3.21 (0.34)** | **3.13 (0.16)** |
|  | e (SE) | 1.13 (0.04) | 1.13 (0.04)) |
|  | Region (SE) |  | 0.71 (2.13) |
| Total variance |  | 32.11 | 32.11 |
| $(a^2 + c^2 + e^2 (+ \text{region}^2))$ | | | |
| A (standardized) |  | 0.640 | 0.640 |
| C (standardized) |  | **0.320** | **0.305** |
| E (standardized) |  | 0.040 | 0.040 |
| REGION (standardized) |  |  | 0.016 |

**Table IV.**
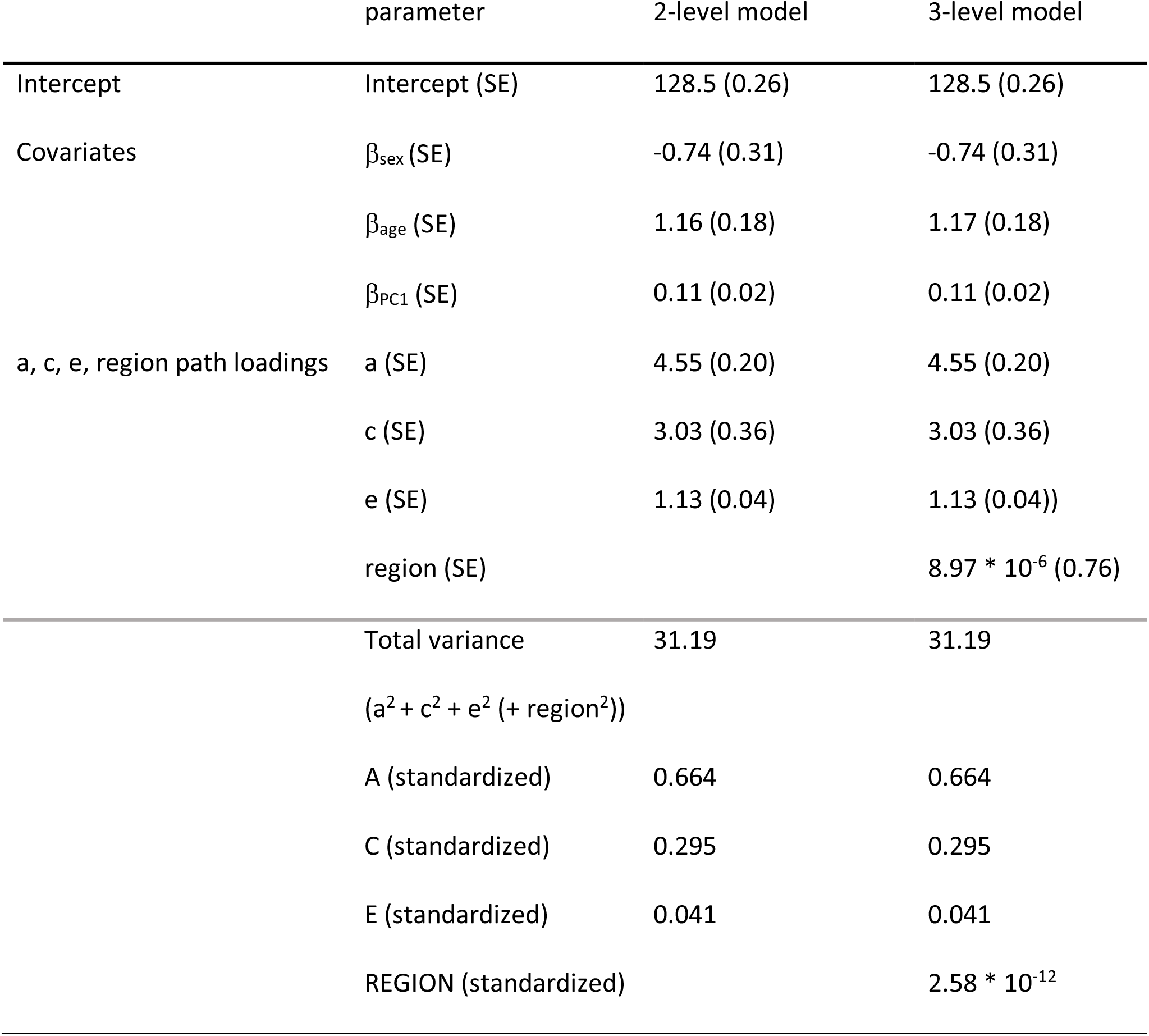
Results of CTM and 2-level and 3-level MLM analyses for the genotyped sample with PC covariate (N = 1375 twins in 714 families). Path coefficient estimates with standard errors (SE) and standardized variance components of the 2- and 3-level ACER model, including PC1.

### Results of analyses of genotyped sample with PC1 as covariate

Table IV displays the parameter estimates and standardized variance components of the model with PC1 as an additional fixed covariate. In the genotyped group, the 2-level model with PC1 as a covariate fitted equally well as the 3-level model with PC1 as a covariate and with region as a third level variable (Δ-2LL < 0.001, Δ df = 1). In this model, the variance attributable to region is zero. So, when PC1 is included as a covariate, region in the 3-level model doesn’t explain any variance above and beyond what is already explained by the first PC in the 2-level model.

## Discussion

In this paper we specified a multilevel twin model in OpenMx and fitted it to data on children’s height. We added a higher-level variable, region in the Netherlands, in which the twin pairs were nested. Adding a third level variable enabled us to address an empirical question, namely whether part of the variance in children’s height can be explained by differences in geographical region.

We found that 68% of the variance in 7-year-old children’s height is attributable to additive genetic factors. Common environmental factors accounted for 26%, and unshared environmental factors for 6% of the variance. We found that regional differences accounted for a significant 1.8% of the phenotypic variance. In a standard multilevel ACE-twin model, ignoring regional clustering, this variance was captured by the C-component. This is expected, because the so called common environmental component captures between-family variance. At age 7, both MZ and DZ twins share region, so that the effect of region will manifest as C.

In a subsample of children who were genotyped and for whom genetic PCs were obtained, we found a statistically significant correlation (*r* = 0.16) between height and the first genetic PC, representing the geographical north-south gradient in the Netherlands. This correlation is similar to previous results for height in a Dutch sample of adults (Abdellaoui et al., 2013). The correlations between the second and third PC and children’s height were negligible. After the inclusion of the first PC in the multilevel analyses, region no longer explained any variance.

This last result indicates that the variance in children’s height that is explained by region is attributable to differences in genetic ancestry. That is, although unmodeled regional clustering manifests as C, it does not mean that the inflation of the common environmental variance is due to genuine shared environmental factors like region. When we included the first PC, which reflects differences in allele frequencies between regions, no variance was explained by geographical region above and beyond what was already explained by the PC. This indicates that a proportion of the variance that is captured in the C-component of the CTM is actually of a genetic nature.

We note some limitations of our study. First, we did not explicitly model qualitative differences in genetic architecture between boys and girls. In the literature there is some evidence that the additive genetic correlation in opposite-sex twins is lower than 0.50, suggesting that partly different genes operate in 7-year-old boys and girls (Silventoinen et al., 2007). However, the twin correlations in our sample did not suggest the presence of qualitative sex differences (we observed correlations of .61 and .63 in the DZ male and DZ female, versus .58 and .68 in the DZ opposite sex male-female and DZ opposite sex female-male twins, respectively).

Secondly, we surmise that the power to detect the region effect in the genotyped sample was low, given the sample size (*N*=1,375 in the genotyped sample). However, the effect sizes in both samples were very similar, and in the full sample (*N*=7,346) the significance of the effect was well established. Therefore, we trust that the regional effect is real.

A final limitation to note is that the current approach assumes that lower levels are fully nested in the higher-level. That is, members of a twin pair cannot differ on the clustering variable. It is therefore not possible to define a third-level clustering variable, when the variable of interest differs within a twin pair (e.g. adult twins who do not live in the same region). It is possible, however, to include variables in which both twins are not nested as a lower-level variance component. When the clustering variable is not specified as a higher-level (i.e., nesting) variable, the effect of clustering can also be manifested as any of the other variance components (i.e., A/C/D/E) when unmodeled. Furthermore, missing data for higher-level clustering variable (here: region) is not allowed. The higher-level variable needs to have a sufficient number of units for the model to have enough power to detect the effect of the higher-level variable (e.g., postal codes in our region example; Goldstein, 2011).

The current study showed that when data are nested in a higher-level variable, adding this higher-level variable to a multilevel model for twin data provides opportunities to further disentangle the etiology of a trait. Clustering can be due to unwanted confounding, for example, batch effects. Applying a multilevel model to identify the nuisance variance that is explained by higher-level clustering would in this case serve as a correction. However, as is shown within this paper, the MLM can also be used to empirically study clustering.

## Acknowledgements and funding

This project is part of the Consortium on Individual Development (CID). CID is funded through the Gravitation Program of the Dutch Ministry of Education, Culture, and Science and the Netherlands Organization for Scientific Research (NWO: 024-001-003). The Netherlands Twin Registry (NTR) is funded by ‘Netherlands Twin Registry Repository: researching the interplay between genome and environment’ (NWO: 480-15-001/674); ‘Twin-family study of individual differences in school achievement’ (NWO: 056-32-010); ‘Longitudinal data collection from teachers of Dutch twins and their siblings’ (NWO: 481-08-011) and BBMRI-NL. EvB is funded by ‘Decoding the gene-environment interplay of reading ability’ (NWO: 451-15-017). MCN, CVD and DIB are funded by NIH grant DA-49867 and MCN previously by DA-018673.

## Compliance with Ethical standards

## Ethical approval

All procedures performed in studies involving human participants were in accordance with the ethical standards of the institutional and/or national research committee and with the 1964 Helsinki declaration and its later amendments or comparable ethical standards.

## Informed consent

Informed consent was obtained from all individual participants included in the study.

